## Supporting Information for "Parameter inference for stochastic biochemical models from perturbation experiments parallelised at the single cell level"

#### A Moment closure

Since the CME is difficult to solve in most cases, a widely used approach is to derive moment equations from it. For mass action kinetics and at most elementary (up to order 2) chemical reactions, these equations can be written as

$$\dot{\eta}(t) = A(\theta)\eta(t) + B(\theta)\bar{\eta}(t), \tag{A.1}$$

where $\eta(t)$ is a column vector containing all uncentered moments up to some order $L$ of $X(t)$, and $\bar{\eta}(t)$ contains uncentered moments of order $L+1$ of $X(t)$. If the reaction network can be perturbed by an external time-varying input, $u(t)$, for instance time varying rates of protein production created by external optical or chemical signals that affect gene expression, these inputs will typically enter the propensity functions, $a_k(X, \theta)$, and the matrices $A(\theta) = A(\theta, t)$ and/or $B(\theta) = B(\theta, t)$ in (A.1) will become explicitly time dependent. If the reaction network is governed by kinetics other than mass action, moment equations can still be derived but the time evolution of the moments $\eta(t)$ in (A.1) will typically depend on moments of order higher than $L+1$. In any case, however, (A.1) cannot be solved exactly due to the dependence of $\dot{\eta}(t)$ on the higher order moments $\bar{\eta}(t)$.

As a consequence, numerous approaches have been proposed to replace the true moment equations by approximate systems

$$\dot{\tilde{\eta}}(t) = f(\tilde{\eta}(t), \theta), \tag{A.2}$$

where the function $f$ can be obtained using approaches such as the linear noise approximation or by replacing the higher order moments with some function of the lower order moments $\bar{\eta} = g(\eta)$ (moment closure), for instance by assuming a specific underlying distribution for $X(t)$ such as a log-normal [31] or Gaussian [34] distribution. How accurate such an approximation is depends on the specific reaction network under consideration and has been studied in numerous cases [4, 10]. In general, however, approximations do often fail and it is *a priori* often hard to say how accurate a given approximation will be.

### B CcaS/CcaR system

#### B.1 The model

We focus on a parametrisation of the stochastic model of the CcaS/CcaR optogenetic system at the single cell level. It is a system where light is used to control molecular biological processes in bacteria. In this system, green light induces the expression of the promoter, the fluorescent protein is produced, hence we can measure fluorescence levels and "count" the fluorescent proteins; while red light stops the production of the protein (see Figure 1 in the main text). We use this system to answer the following questions: For a fixed number of cells in the experiment is there a benefit of diversifying light stimulation patterns at the single cell level? How many cells need to be observed to allow for accurate inference of model parameters?

We perform parameter inference for the model proposed in Chait et al. (2017). It is a stochastic single cell model given by the following reaction network:

$$\emptyset \xrightarrow{h_0} E \xrightarrow{h} \emptyset, \qquad \emptyset \xrightarrow{aE(t)L(t)} F \xrightarrow{b} \emptyset, \tag{B.3}$$

where $L(t) = \frac{(cl(t))^n}{(cl(t))^n + k^n}$, and the dynamics of $l(t)$ is given by:

$$\frac{dl(t)}{dt} = u(t) - cl(t). \tag{B.4}$$

The stochastic variable $E(t)$ is called *responsiveness* of the cell. It represents a fluctuating state of a cell and it captures cell-to-cell variability in numbers of ribosomes, plasmid copy numbers *etc.* It is assumed to be a stationary Poisson process that can be fully characterised by its mean $m := \frac{h_0}{h}$ and its time scale $h$. It is a hidden variable, meaning we do not measure it directly, but we estimate its state and its parameters through the model fitting.

The second stochastic variable $F(t)$ represents the *number of fluorescent proteins* in a cell and is measured in experiments. The actual measured quantity is fluorescence $s \cdot F(t)$ with noise as defined in equation (3) (in the main text), where $s$ is a proportionality constant. The state of $F(t)$ depends on the hidden variable $E(t)$, *i.e.* on the state of the cell, and on the deterministic but non-linear variable $L(t)$ that captures *the state of the light system*. The variable $L(t)$ is given as a Hill function that depends on the deterministic *light–activation variable* $l(t)$, which on the other hand depends on a *light signal* $u(t)$. The light signal function, $u(t)$, is a step function that switches between $u(t) = 1$, for green light and $u(t) = 0$, for red light. Measurements are taken every $t_m = 6\,\text{min}$. Finally, parameter inference is used to identify 8 parameters, $\theta = \{a, b, s, m, h, c, n, k\}$, from data collected from single cells.

#### B.2 Simulated single cell data

Simulation of the model (B.3)-(B.4) was implemented in MATLAB using the parameters from Table B.1. These parameters are chosen in order to obtain system dynamics that are similar to experimentally observed ones.

**Table B.1:** Exact values of parameters of the model (11)-(13)

| parameter | $a$ | $b$ | $s$ | $m$ | $h$ | $c$ | $n$ | $k$ |
|---|---|---|---|---|---|---|---|---|
| exact value | 0.6228 | 0.0113 | 1.5631 | 4.1176 | 0.0059 | 0.0842 | 1.4312 | 0.5818 |

Using Gillespie's algorithm, we generated data sets with different numbers of cells per data set (12, 30 or 60) and with different pre-defined light inputs (4 groups where all cells receive the same light input and one group where cells get one of 6 different light inputs) as explained in Section 3.2.1. The simulated data are shown in Figures B.1 - B.3 .

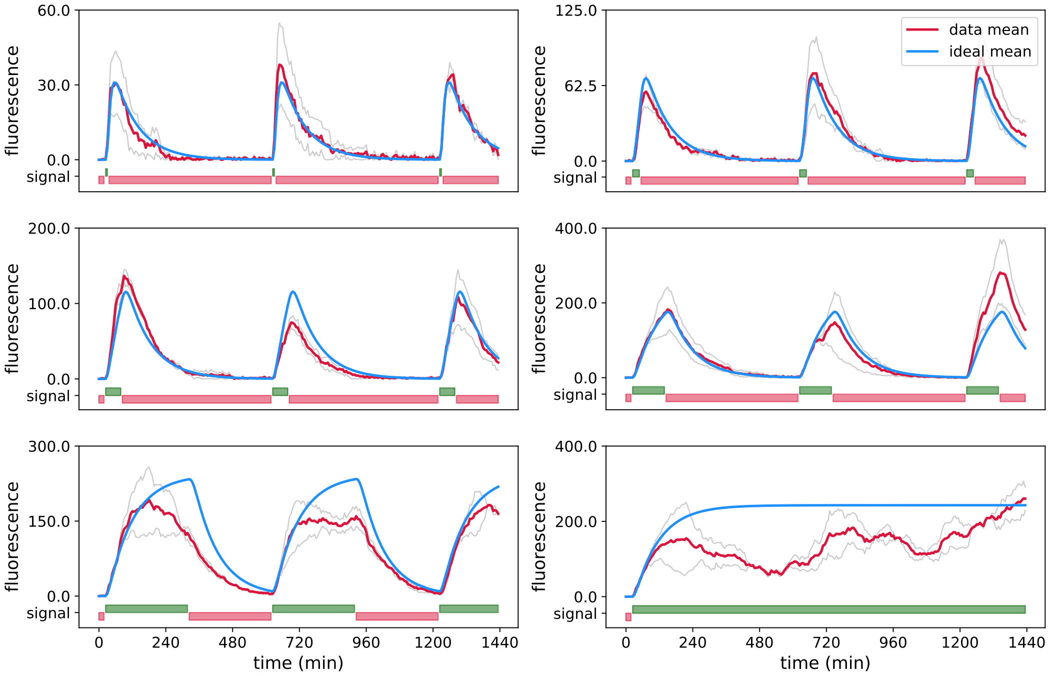


**(a)** Data: G0, 12 cells in total, 2 cells per light group.

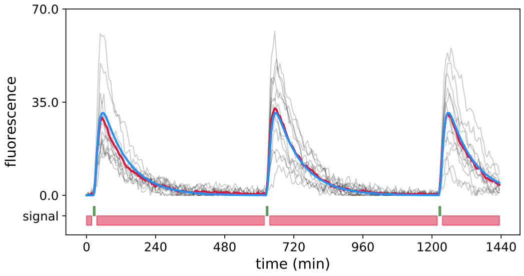


**(b)** Data: G1, 12 cells.

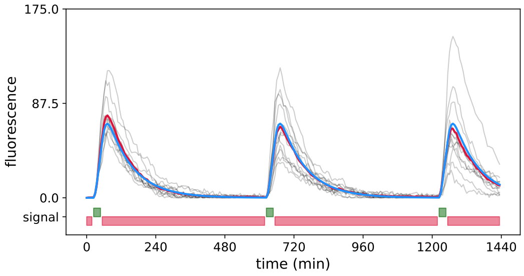


**(c)** Data: G2, 12 cells.

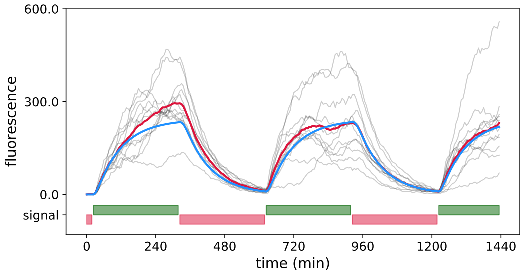


**(d)** Data: G5, 12 cells.

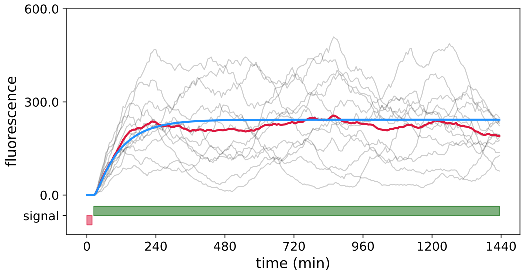


**(e)** Data: G6, 12 cells.

**Figure B.1:** Simulated data using Gillespie's algorithm for 12 cells per data set. In each panel gray lines represent simulated trajectories, red lines are means of simulated data, blue lines are theoretical means, obtained with the model and the exact parameters used for data simulation. Red and green light inputs are shown in red and green bars, respectively, at the bottom of each panel. Note that each light pattern starts with red light.

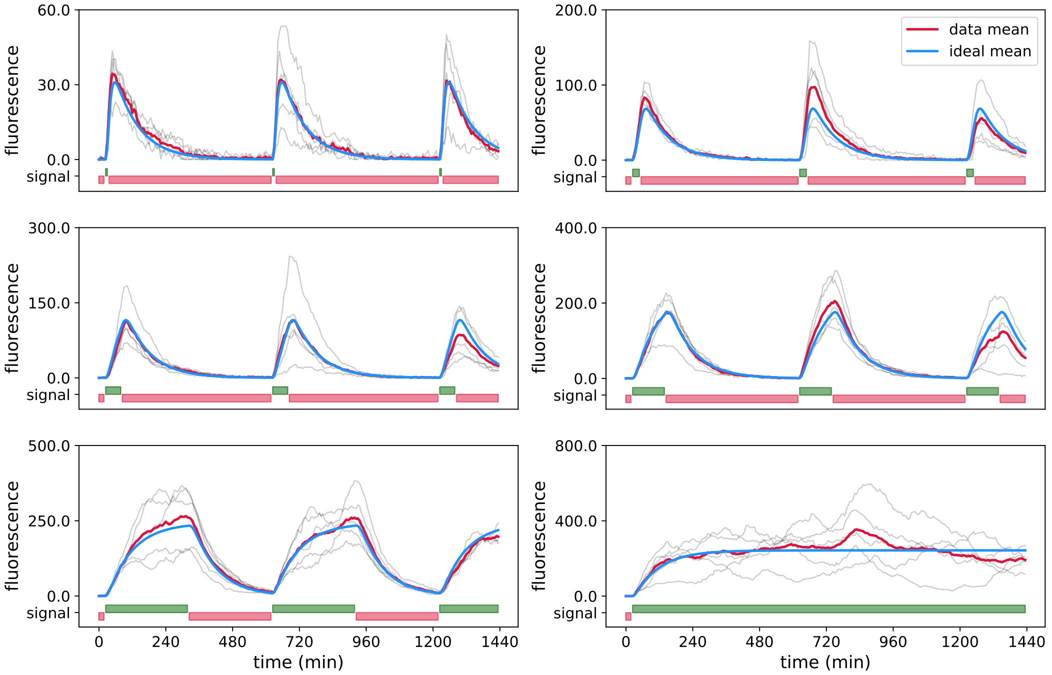


**(a)** Data: G0, 30 cells in total, 5 cells per light group.

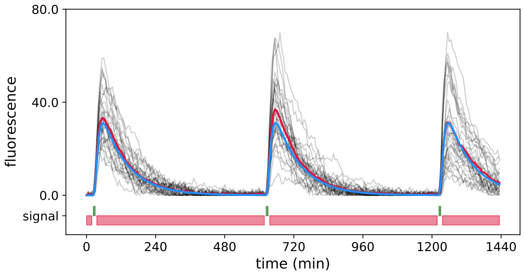


**(b)** Data: G1, 30 cells.

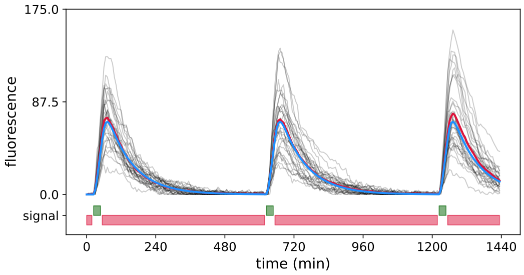


**(c)** Data: G2, 30 cells.

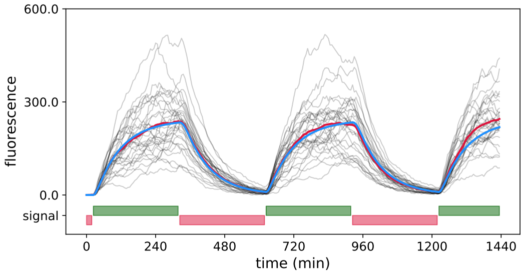


**(d)** Data: G5, 30 cells.

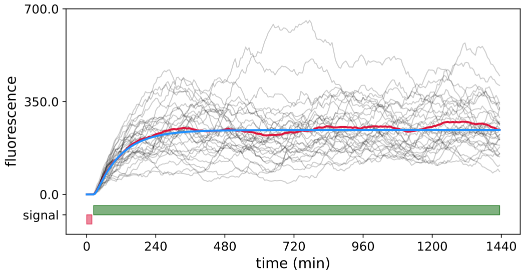


**(e)** Data: G6, 30 cells.

**Figure B.2:** Simulated data using Gillespie's algorithm for 30 cells per data set. Colour coding is equivalent to Figure B.1.

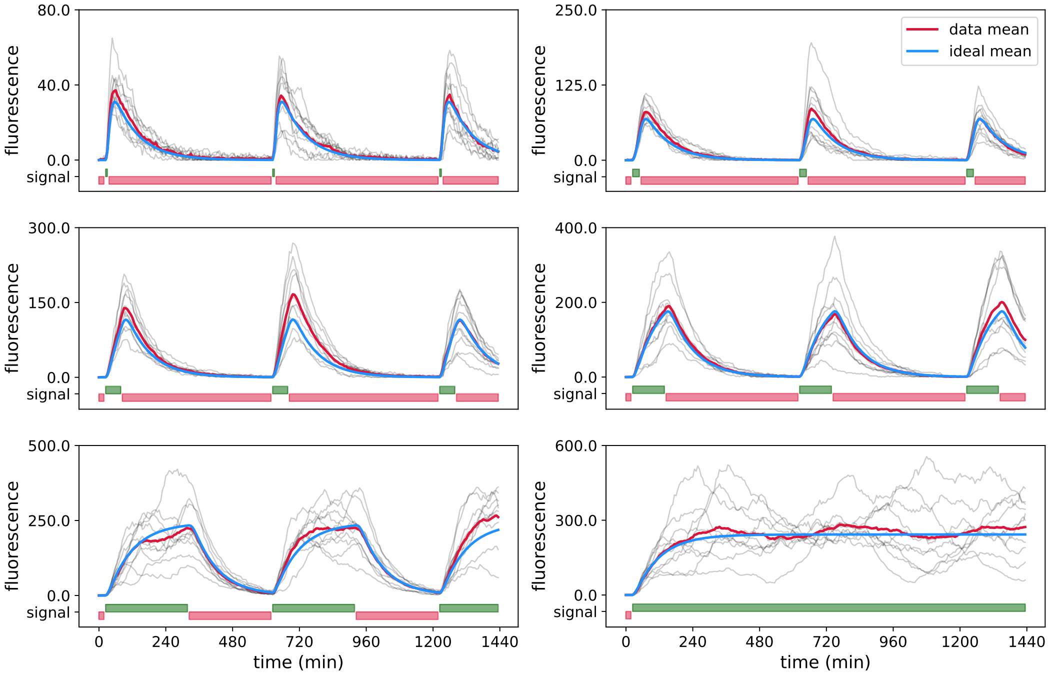


**(a)** Data: G0, 60 cells in total, 10 cells per light group.

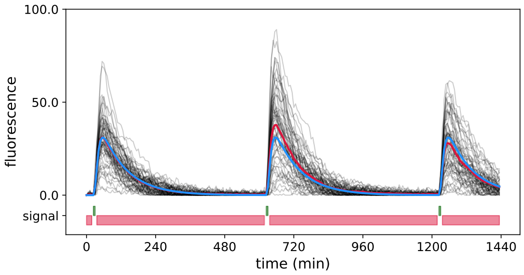


**(b)** Data: G1, 60 cells.

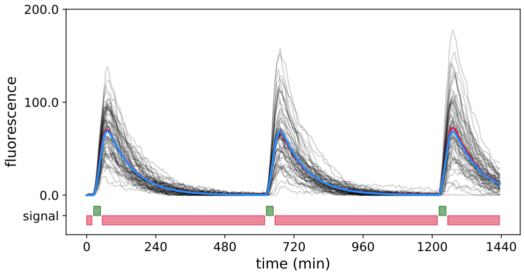


**(c)** Data: G2, 60 cells.

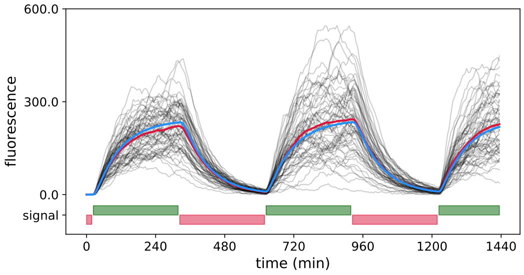


**(d)** Data: G5, 60 cells.

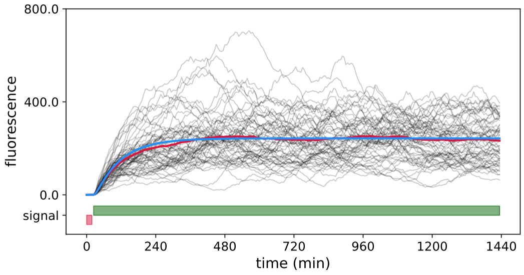


**(e)** Data: G6, 60 cells.

**Figure B.3:** Simulated data using Gillespie's algorithm for 60 cells per data set. Colour coding is equivalent to Figure B.1.

#### B.3 Implementation of LNA with Kalman filter

In order to compute the likelihood of data for a given set of parameters of the model given in (B.3) - (B.4), we implemented Algorithm 1, in MATLAB, as follows. We create a function that takes as input:

- $n$ cell trajectories, $y^{all}$, where each trajectory is given as an array $y = (y_0, y_1, y_2, ...., y_S)$ where $y_i \in \mathbb{R}^+$;
- a light input $u$ for each trajectory $y$, which is an array of 0s and 1s which correspond to red and green light signals, respectively, *i.e.* $u = (u_0, u_1, u_2, ...., u_S)$ where $u_i \in \{0, 1\}$;
- a set of parameters $\theta = \{a, b, s, m, h, c, n, k\}$ for which we want to compute the likelihood.

For each trajectory $y$ and its corresponding light input $u$ we implement the following steps:

1. Solve the ODE (B.4), using light input $u$ and parameters $c, n$ and $k$, and compute and save values of $L(t)$ for each time step of the simulation.
2. Every simulated trajectory starts from the fixed state, hence the moments for the initial states are also fixed to $\mu_0 = (m\ 0)$ and $\Sigma_0 = \begin{pmatrix} 0 & 0 \\ 0 & 0 \end{pmatrix}$, and the log-likelihood is $\log(p(y_0 \mid \theta)) = 0$.
3. Iterate the following steps for measurements $y_i, i = 1, ..., S$ of the given trajectory $y$:
   (a) Use Kalman filter with $\mu_{i-1}, \Sigma_{i-1}, y_{i-1}$ and noise parameter $\sigma$, to update estimates $\mu_{i-1}$ and $\Sigma_{i-1}$. For $R = \sigma^2$ and $C = (0\ s)$, the classical Kalman update is given as

$$K = \Sigma_{i-1} C^T (C \Sigma_{i-1} C^T + R)^{-1},$$
$$\mu^*_{i-1} = \mu_{i-1} + K(y_{i-1} - \mu_{i-1} C^T),$$
$$\Sigma^*_{i-1} = (I - KC)\Sigma_{i-1}.$$

(b) From updated estimates of centered moments

$\mu^*_{i-1} = (\mu^*_E \;\; \mu^*_F)$ and $\Sigma^*_{i-1} = \begin{pmatrix} \mu^*_{EE} & \mu^*_{EF} \\ \mu^*_{EF} & \mu^*_{FF} \end{pmatrix}$,

calculate the uncentered moments

$\overline{\mu}^*_{i-1} = (\mu^*_E \;\; \mu^*_F)$ and $\overline{\Sigma}^*_{i-1} = \begin{pmatrix} \mu^*_{EE} + \mu^{*2}_E & \mu^*_{EF} + \mu^*_E\mu^*_F \\ \mu^*_{EF} + \mu^*_E\mu^*_F & \mu^*_{FF} + \mu^{*2}_F \end{pmatrix}$,

and use them as initial conditions for the system of ODEs of the moment equations on the time interval $[t_{i-1}, t_i]$. Using pre-calculated values for function $L(t)$, solve:

$$\begin{aligned}
\frac{\mathrm{d}\mu_E}{\mathrm{d}t} &= mh - \mu_E h, \\
\frac{\mathrm{d}\mu_F}{\mathrm{d}t} &= aL(t)\mu_E - b\mu_F, \\
\frac{\mathrm{d}\mu_{EE}}{\mathrm{d}t} &= mh + 2mh\mu_E + h\mu_E - 2h\mu_{EE}, \\
\frac{\mathrm{d}\mu_{EF}}{\mathrm{d}t} &= mh\mu_F - h\mu_{EF} - b\mu_{EF} + aL(t)\mu_{EE}, \\
\frac{\mathrm{d}\mu_{FF}}{\mathrm{d}t} &= b\mu_F - 2b\mu_{FF} + aL(t)\mu_E + 2aL(t)\mu_{EF}.
\end{aligned}$$

Obtain estimates of the uncentered moments at time $t_i$ as

$\overline{\mu}_i = (\mu_E \;\; \mu_F)$ and $\overline{\Sigma}_i = \begin{pmatrix} \mu_{EE} & \mu_{EF} \\ \mu_{EF} & \mu_{FF} \end{pmatrix}$.

The centered moments, that are used in Kalman filter, calculate as

$\mu_i = (\mu_E \;\; \mu_F)$ and $\Sigma_i = \begin{pmatrix} \mu_{EE} - \mu^2_E & \mu_{EF} - \mu_E\mu_F \\ \mu_{EF} - \mu_E\mu_F & \mu_{FF} - \mu^2_F \end{pmatrix}$.

(c) Update log-likelihood using $\Sigma_{FF} = s^2(\mu_{FF} - \mu^2_F) + \sigma^2$ and the observation $y_i$,

$$\log(p(y_i, ..., y_0 \mid \theta)) = \log(p(y_{i-1}, ..., y_0 \mid \theta)) - \frac{1}{2}\left(\log\left(2\pi\Sigma_{FF}\right) + \frac{(s\mu_F - y_i)^2}{\Sigma_{FF}}\right).$$

#### B.4 Inference of parameters using Metropolis Hastings MCMC

To infer a model parameter from simulated or experimental data, we need to search for the value of that parameter that best describe the observed data. We might be interested in the exact value with the highest likelihood, such as maximum likelihood (MLE) or maximum a posteriori (MAP) estimation, or in the posterior distribution that shows values that are more likely to be true. Depending on the problem and the model, there might be two or more values with similar likelihoods, and in this case the posterior distribution is more informative. On the other hand, when we need a concrete value for further investigations (*i.e.* for concrete predictions) the exact value may be more practical.

In our case, we need to explore an 8-dimensional space in order to infer $\theta = \{a, b, s, m, h, c, n, k\}$. To that end, we use a Metropolis Hastings Markov chain Monte Carlo (MCMC) method as it provides us with an unbiased posterior distribution of the parameters. For each sampled set of parameters $\theta$, we compute the likelihood $p(y \mid u, \theta)$ using the LNA with Kalman filter, as described in the previous section. Further, since all model parameters are positive, we use as proposal distribution $g(\theta_{\text{new}}|\theta_{\text{old}})$ a log-normal distribution with mean $\mu = \log(\theta_{\text{old}})$ and standard deviation $\sigma_g = 0.02 \cdot \mathbb{I}$, where $\mathbb{I}$ is an identity matrix. The value for $\sigma_g$ was obtained empirically in order to get an acceptance rate of the chain around 0.3. Additionally, we reject propositions that do not fall in the limits for the two parameters, $n < 10$ and $k < 1$, and keep drawing a new set of parameters until they do.

One should note that our likelihood evaluation algorithm relies on the assumption that local states of the system follow Gaussian distributions. Hence, when estimating, the algorithm allows negative values of the mean responsiveness $\mu_E$ and mean fluorescence $\mu_F$, which do not make sense and have to be corrected. Similarly, due to approximation errors of the LNA, likelihood calculations may lead to spurious results. If a proposed $\theta_{\text{new}}$ leads to such issues, which are rare for simulated data, we reject it and resample.

We run the search for parameters for each of the data sets given in Figures B.1 - B.3. We run chains of 10 000 iterations and remove 4 000 iterations as burn-in period. In the following, we plot one-dimensional and two-dimensional marginals of posterior distributions comparing results for different data sets. Results shown in Figure 5 and Figures B.4-B.11.

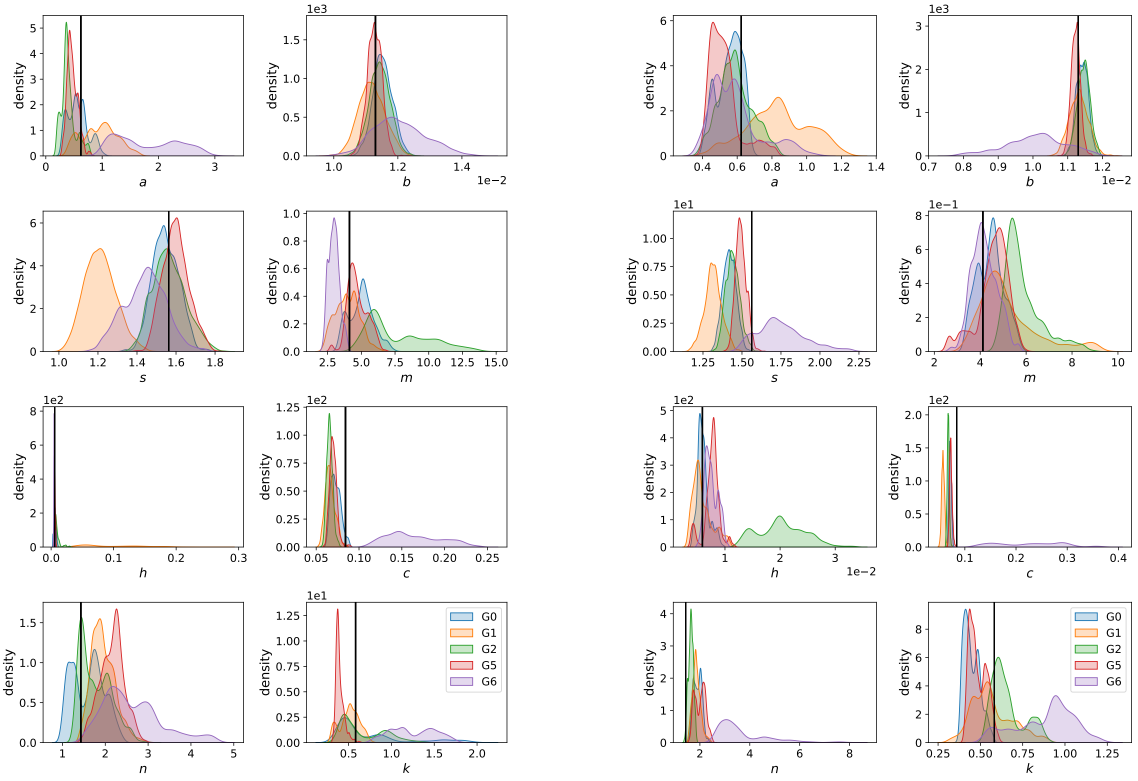


**(a)** posterior distributions for 12 cells

**(b)** posterior distributions for 30 cells

**Figure B.4: Posterior distribution comparison for all groups.** **(a)** One-dimensional marginals of posterior distributions obtained using 12 cells, either all exposed to the same light input (G1 - orange, G2 - green, G5 - red, G6 - purple) or to diversified light inputs containing 2 cells for each of the 6 light patterns (G0 - blue). The vertical black lines in each panel show the true value of the corresponding parameter that was used to simulate the data sets. **(b)** One-dimensional marginals of posterior distributions obtained using 30 cells. Colour coding is the same as in panel (a). In all cases, posterior distributions have been obtained as histograms of values visited by the MCMC search of 10 000 iterations, without a burn-in period of 4 000 iterations.

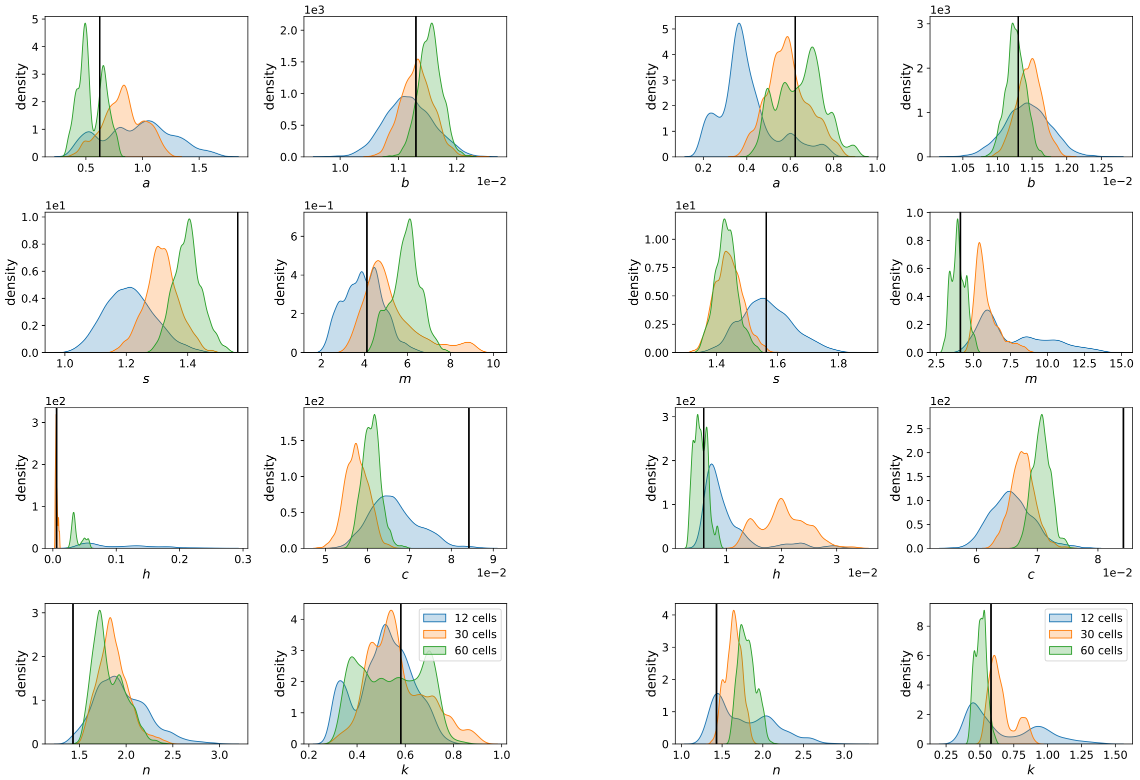


**(a)** posteriors for G1 **(b)** posteriors for G2

**Figure B.5: Posterior distribution comparison for G1 and G2. (a)** One-dimensional marginals of posterior distributions obtained using data of G1, i.e. with all cells exposed to the same light signal, with different numbers of cells (12 - blue, 30 - orange, 60 - green). The vertical black lines in each panel show the true value of the corresponding parameter that was used to simulate the data sets. **(b)** One-dimensional marginals of posterior distributions obtained using data of G2. Colour coding is the same as in panel (a). In all cases, posterior distributions have been obtained as histograms of values visited by the MCMC search of 10 000 iterations, without a burn-in period of 4 000 iterations.

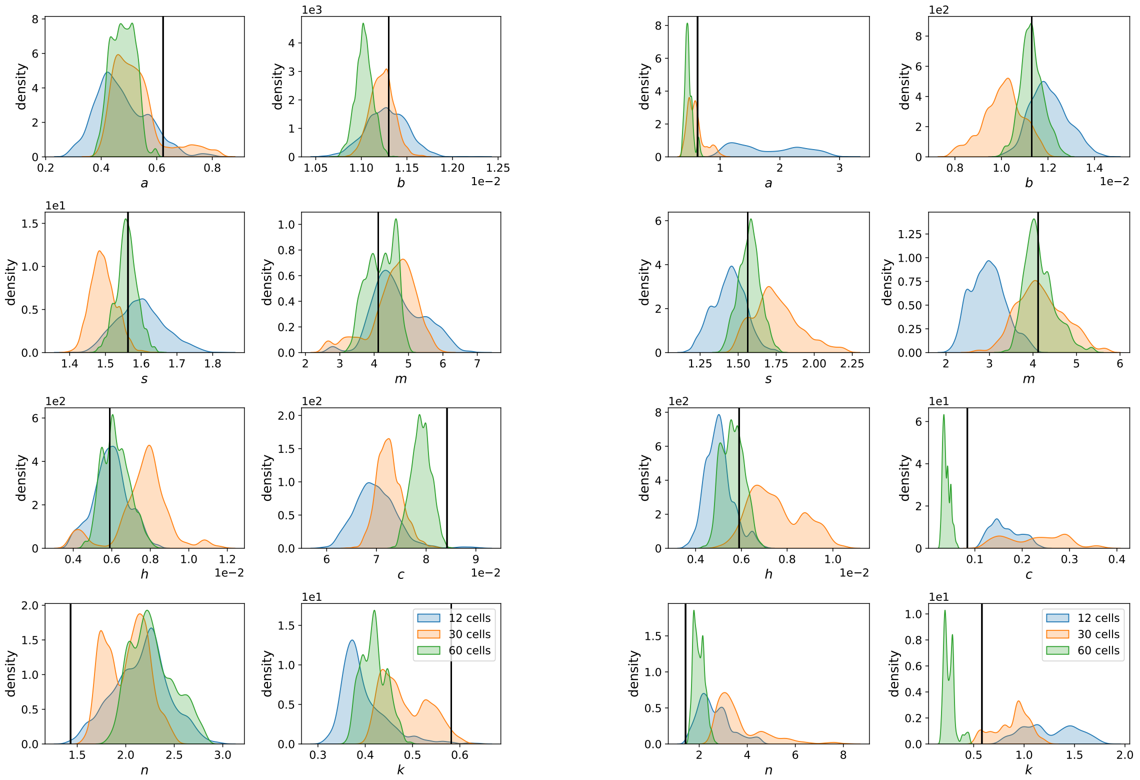


**(a)** posteriors for G5

**(b)** posteriors for G6

**Figure B.6: Posterior distribution comparison for G5 and G6. (a)** One-dimensional marginals of posterior distributions obtained using data of G5, i.e. with all cells exposed to the same light signal, with different numbers of cells (12 - blue, 30 - orange, 60 - green). The vertical black lines in each panel show the true value of the corresponding parameter that was used to simulate the data sets. **(b)** One-dimensional marginals of posterior distributions obtained using data of G6. Colour coding is the same as in panel (a). In all cases, posterior distributions have been obtained as histograms of values visited by the MCMC search of 10 000 iterations, without a burn-in period of 4 000 iterations.

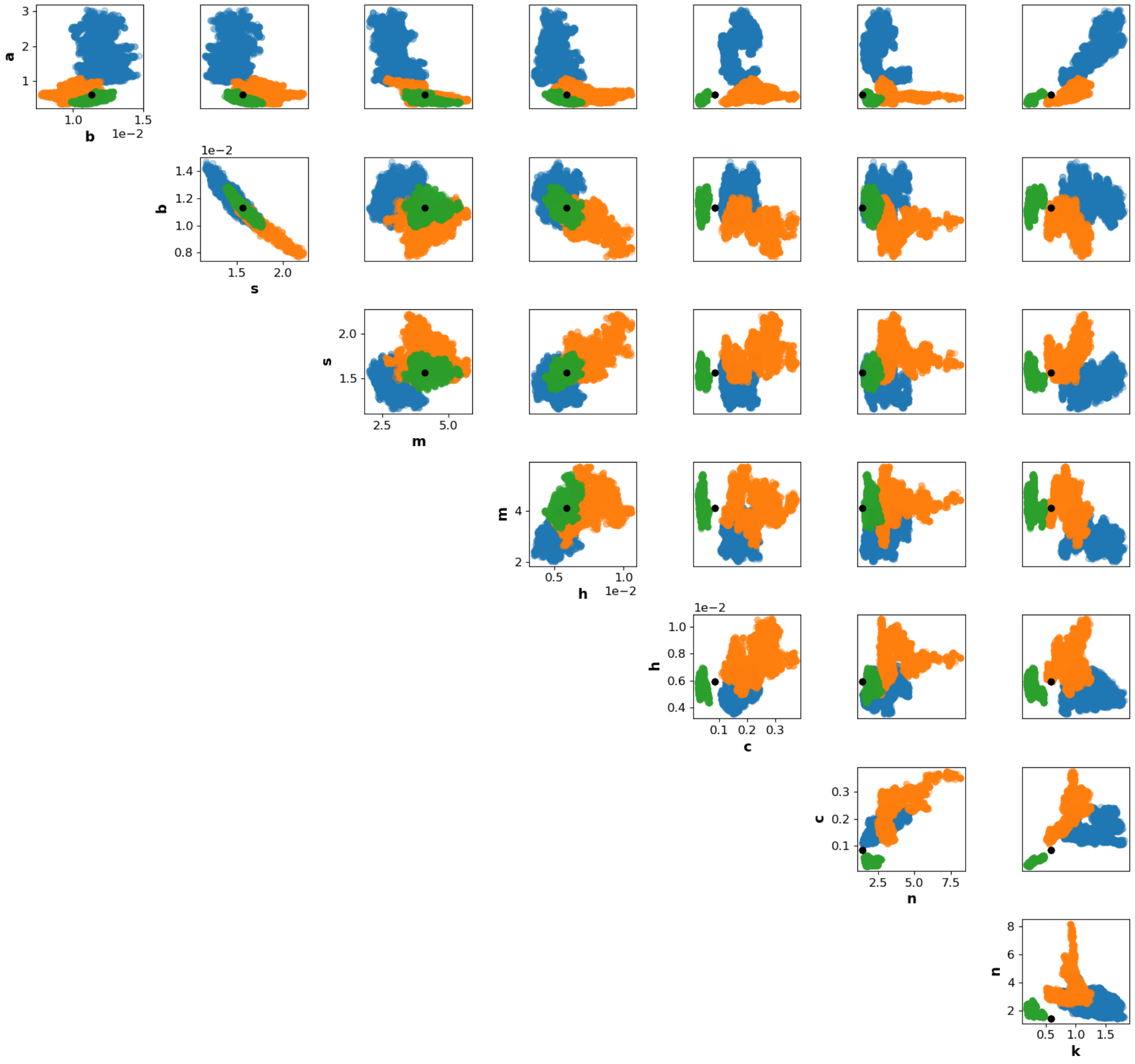


**Figure B.7: Two-dimensional marginals of posterior distributions for G6.** Sets of data with 12 (blue), 30 (orange) and 60 (green) cells. The black dots represent the place of the exact parameters used for data simulations.

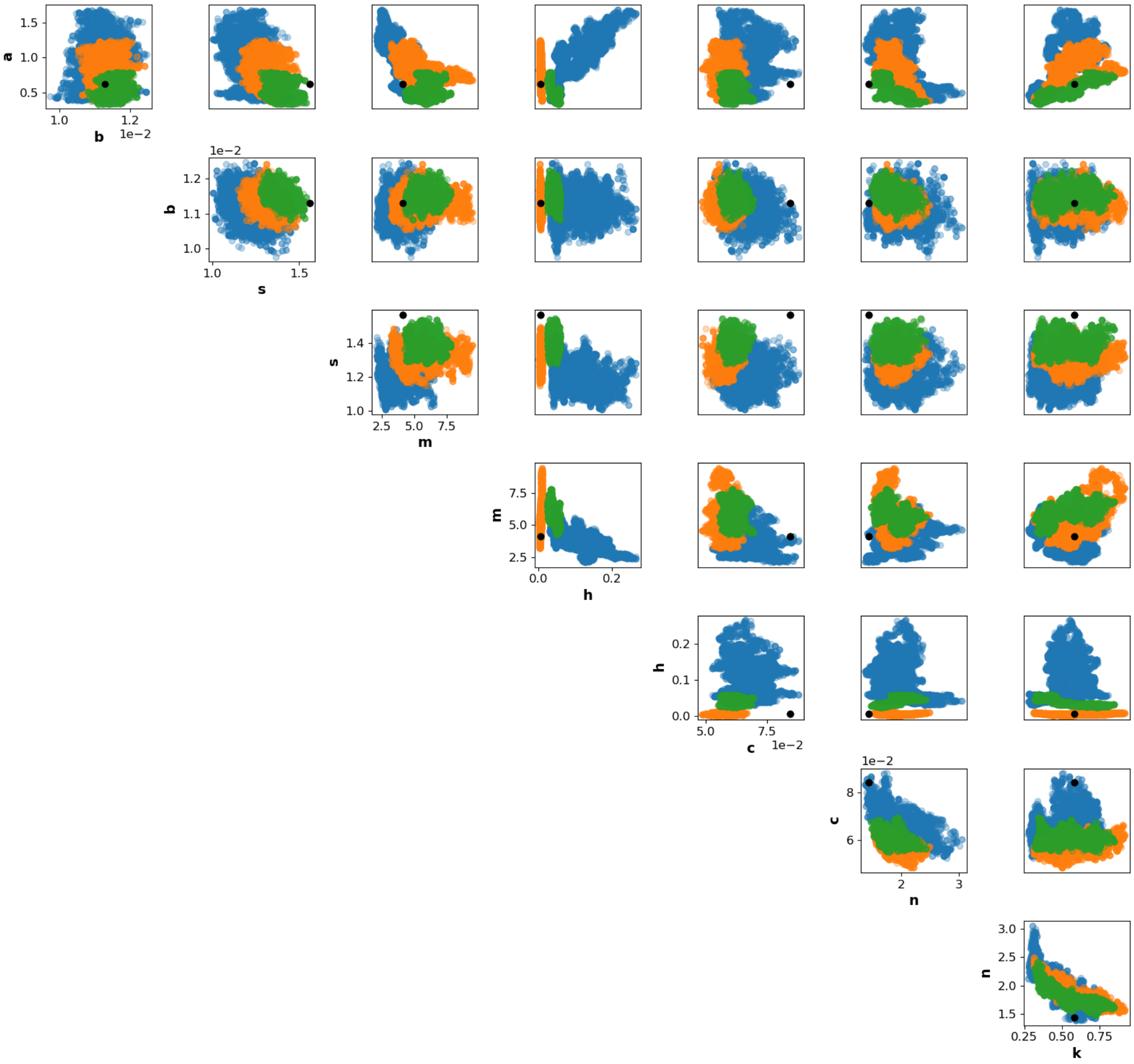


**Figure B.8: Two-dimensional marginals of posterior distributions for G1.** Colour coding is the same as in Figure B.7.

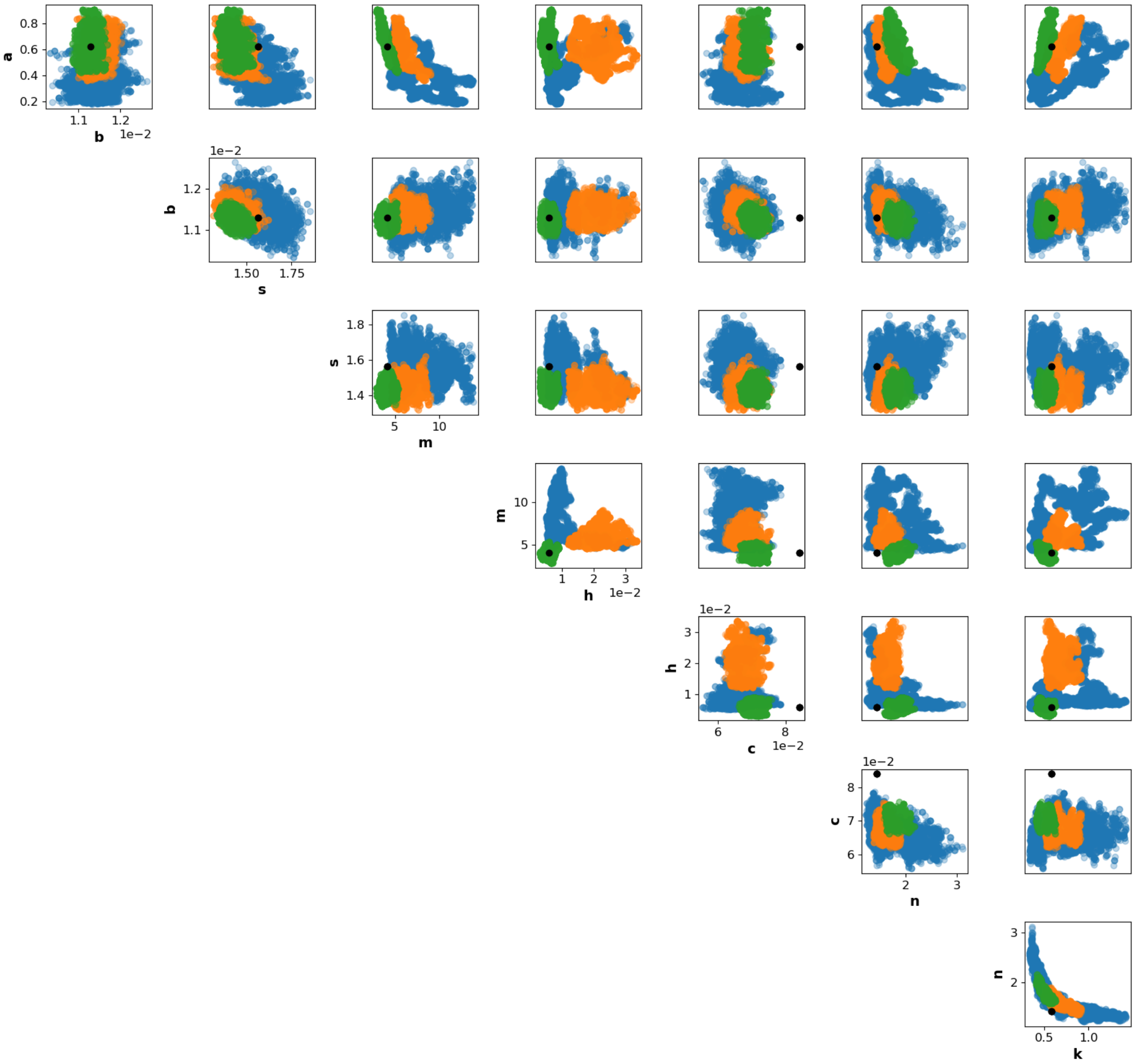


**Figure B.9: Two-dimensional marginals of posterior distributions for G2.** Colour coding is the same as in Figure B.7.

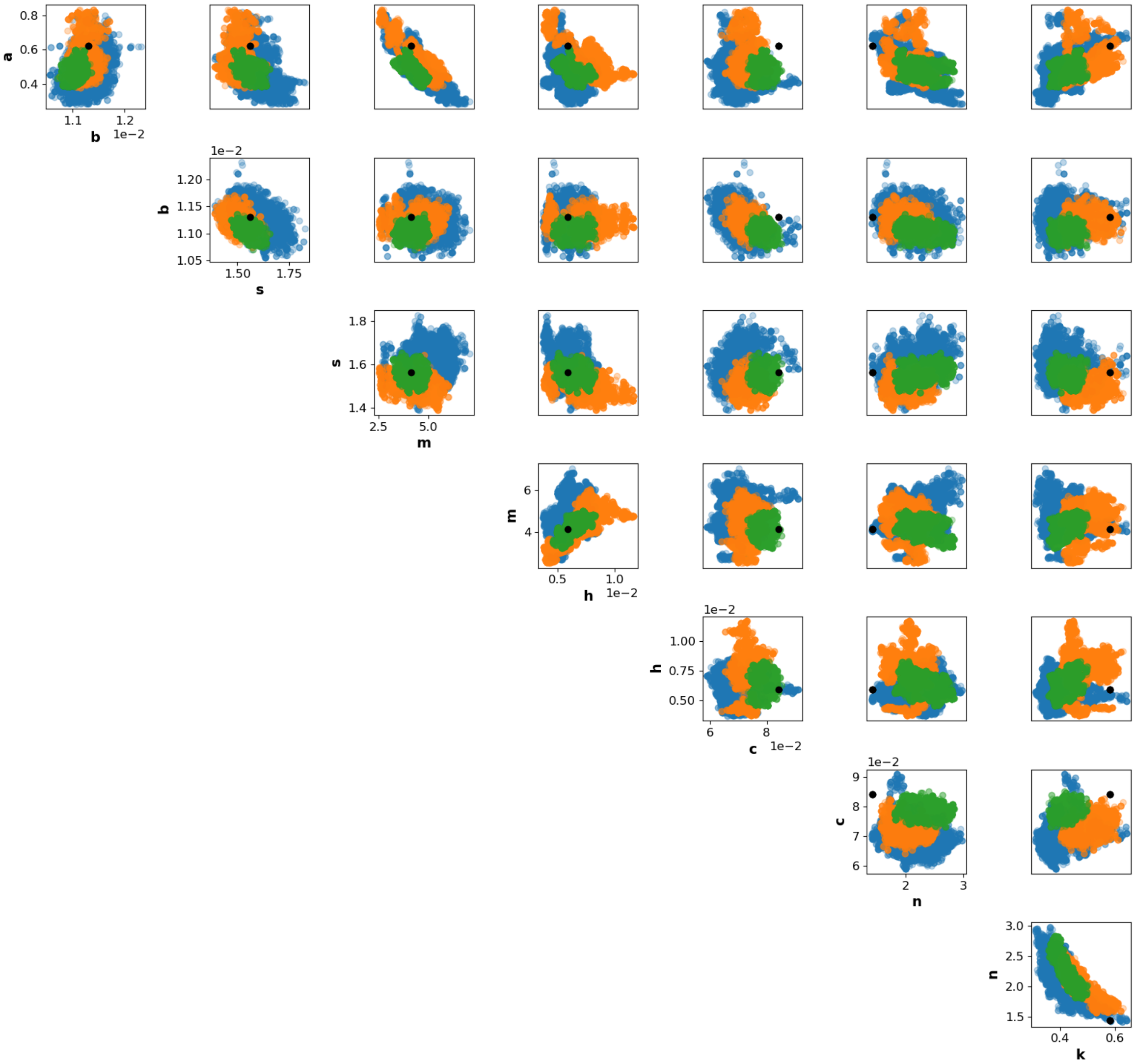


**Figure B.10: Two-dimensional marginals of posterior distributions for G5.** Colour coding is the same as in Figure B.7.

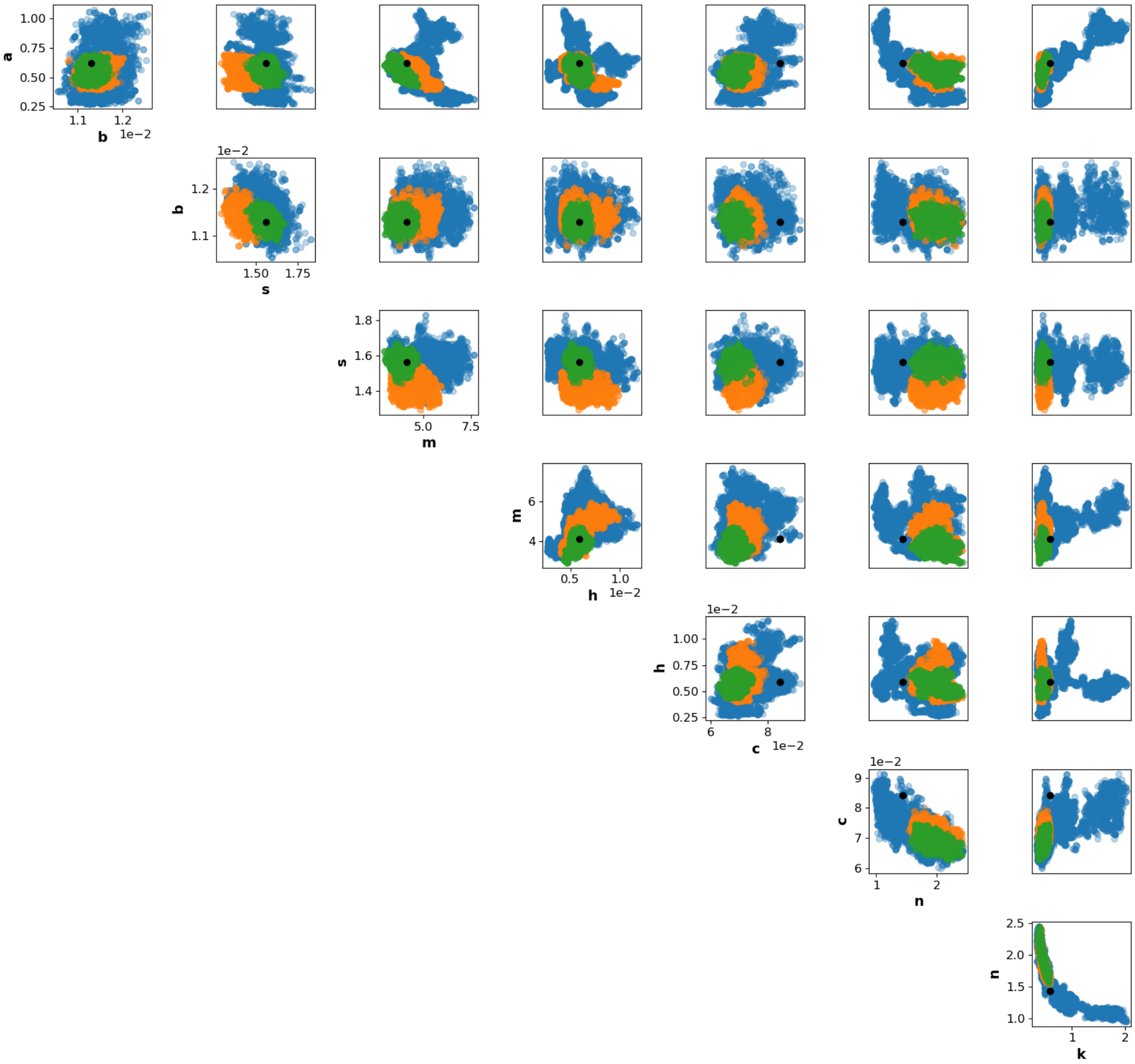


**Figure B.11: Two-dimensional marginals of posterior distributions for G0.** Colour coding is the same as in Figure B.7.

#### B.5 Validation of the method

It is interesting and important for practical purposes to determine how well the outcomes of experiments that have not been performed can be predicted based on the model and parameters that have been learnt from the experiments that were performed. If parameters have been learnt from only one data set, then we can test model predictions on all other data sets so these other data sets can be used as proxies for experiments that have not yet been performed. Since the model is stochastic, comparing single cell trajectories could be misleading. Instead, we compare the mean of observed data to the deterministic mean computed using the model with parameters fixed to the maximum a posteriori estimator (MAP).

For conciseness, we focus only on data with 60 cell trajectories. The values of parameters that we learnt using these simulated data are given in Table B.2.

Firstly, we verify that the model with inferred parameters from a given data set fits well the mean of that same data set. These results are shown in Figure B.12. Then, we investigate how the model with a MAP estimator obtained from one data set predicts the behaviour of another data set. Results are presented in Figure B.13. We find that the model using any of the MAP estimators fits almost any other data quite well, which means that our algorithm is generally well suited for parameter inference in this case. Furthermore, we observe that, like in the previous analyses, the model using the MAP estimator obtained from G1 is clearly worse in predicting the behaviour of data that received signals 3, 4, 5 or 6. Also, if we use the model with the MAP estimator obtained by fitting G6, we do not predict well the behaviour of cells that received signals 1 or 2, but neither of cells that received signals 3, 4 or 5. In this analysis, we do not see a significant difference between predictions resulting from using the MAP estimators obtained from G0, G5 and G2.

**Table B.2:** MAP estimators for parameters of the model (11)-(13) obtained using different data sets where each data set has 60 cell trajectories.

| parameter | $a$ | $b$ | $s$ | $m$ | $h$ | $c$ | $n$ | $k$ |
|---|---|---|---|---|---|---|---|---|
| exact value | 0.6228 | 0.0113 | 1.5631 | 4.1176 | 0.0059 | 0.0842 | 1.4312 | 0.5818 |
| MAP G0 | 0.5583 | 0.0114 | 1.5680 | 3.7287 | 0.0054 | 0.0680 | 1.9014 | 0.4548 |
| MAP G1 | 0.5016 | 0.0116 | 1.3750 | 6.0474 | 0.0352 | 0.0611 | 1.8448 | 0.4881 |
| MAP G2 | 0.5708 | 0.0113 | 1.4359 | 4.3500 | 0.0054 | 0.0700 | 1.8080 | 0.4959 |
| MAP G5 | 0.4494 | 0.0110 | 1.5698 | 4.3230 | 0.0063 | 0.0778 | 2.3618 | 0.4091 |
| MAP G6 | 0.4785 | 0.0113 | 1.5710 | 4.0427 | 0.0053 | 0.0338 | 1.8926 | 0.2283 |

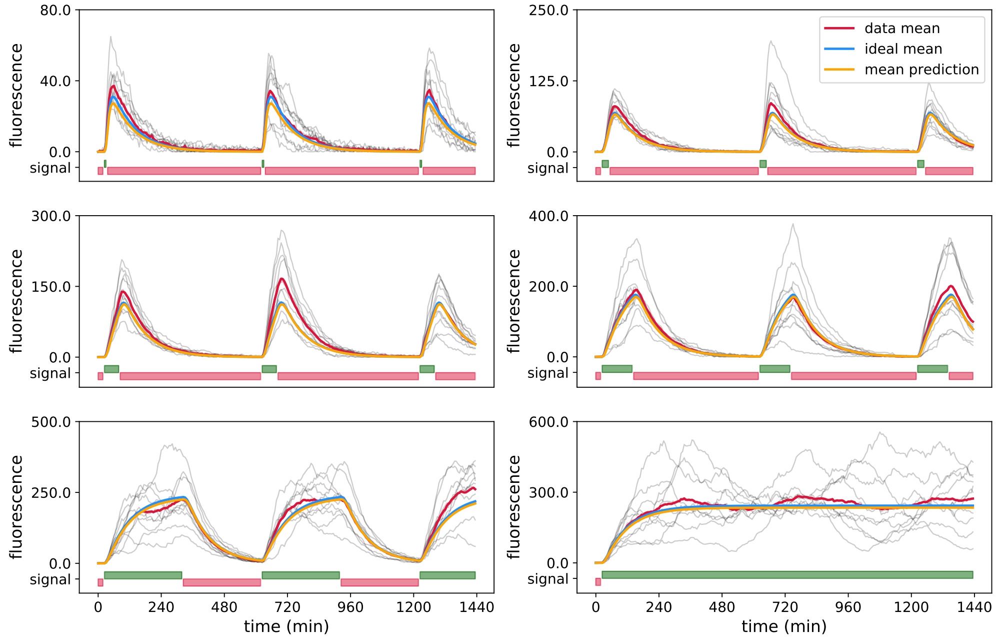


**(a)** Data: G0, 60 cells in total, 10 cells per light group.

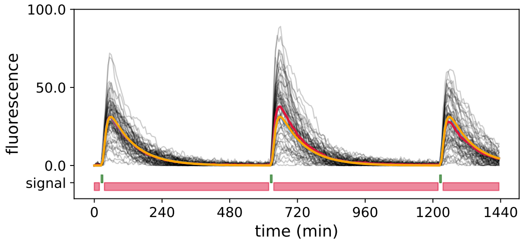


**(b)** Data: G1, 60 cells.

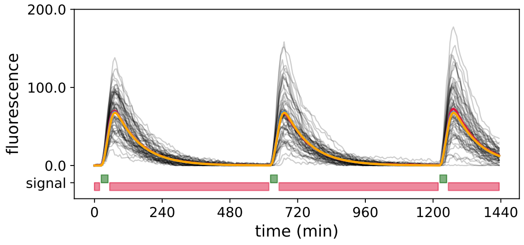


**(c)** Data: G2, 60 cells.

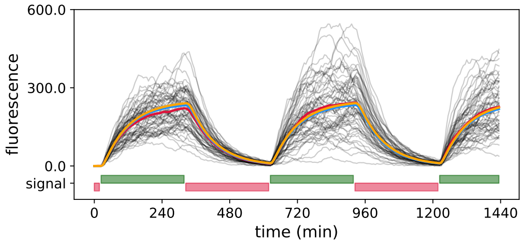


**(d)** Data: G5, 60 cells.

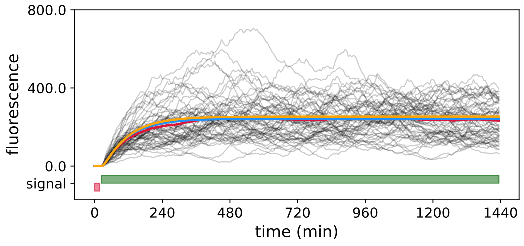


**(e)** Data: G6, 60 cells.

**Figure B.12: Comparing observed mean fluorescence and theoretical mean using the model and MAP estimators of the same data set. (a)-(e)** Each gray line represents one simulated trajectory, red lines are means of simulated data, blue lines are theoretical means of the model with the exact parameters used for simulation and yellow lines are the predicted means using MAP estimators as model parameters. We used data sets with 60 cell trajectories. Red and green light inputs are shown in red and green bars, respectively, at the bottom of each panel. Note that each light pattern starts with red light.

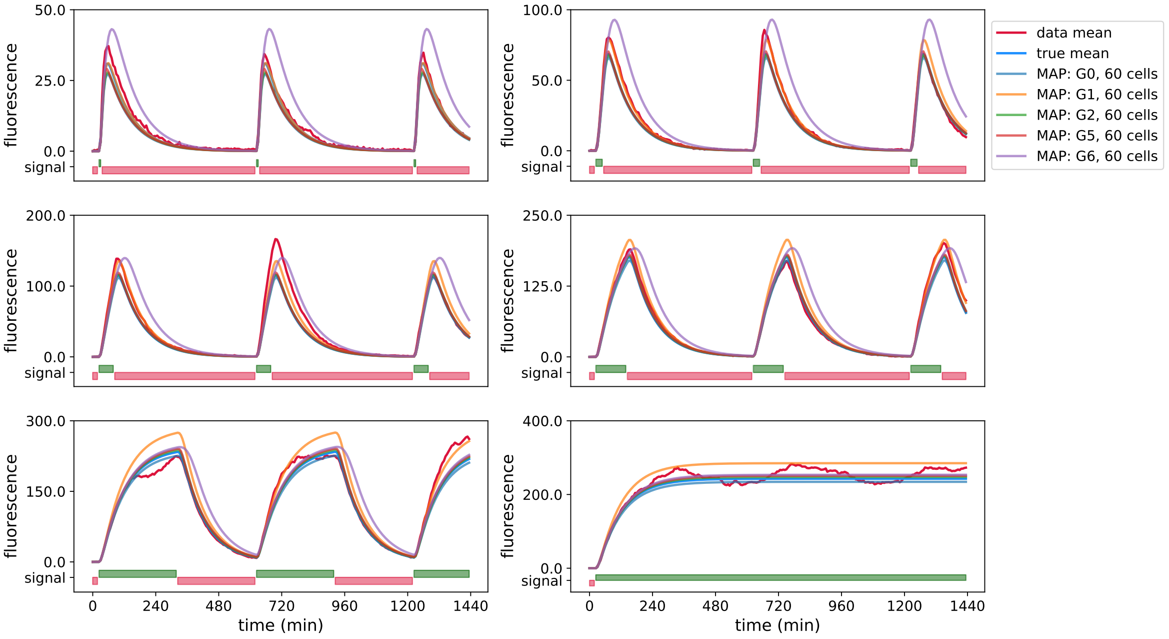


**(a)** Data: G0, 60 cells in total, 10 cells per light group.

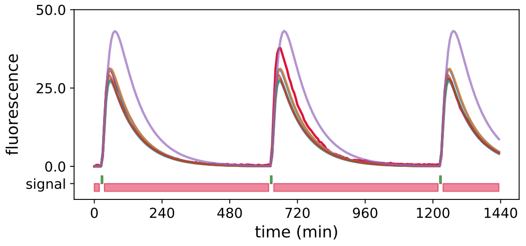


**(b)** Data: G1, 60 cells.

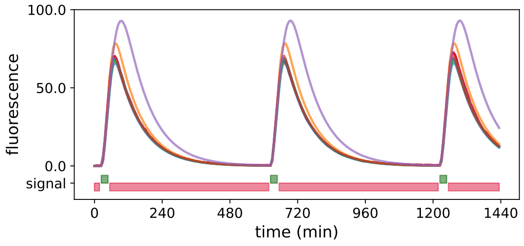


**(c)** Data: G2, 60 cells.

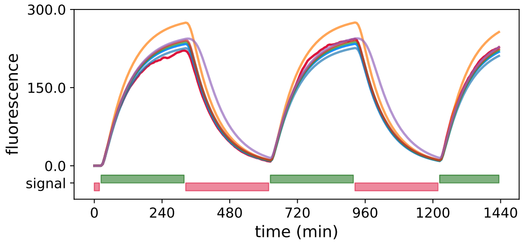


**(d)** Data: G5, 60 cells.

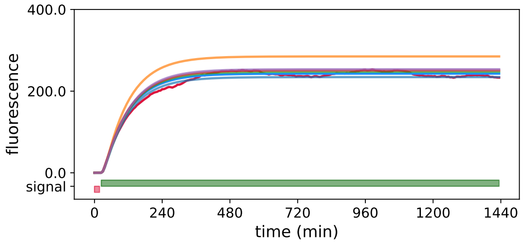


**(e)** Data: G6, 60 cells.

**Figure B.13: Predicting mean fluorescence levels using the model and MAP estimators of different data sets. (a)-(e)** Red lines are means of simulated data, blue lines are theoretical means of the model with the exact parameters used for simulation and other lines are theoretical means obtained with the model and MAP estimators of different data sets. Green lines represent the light input, where each pattern starts with a red light signal. We used data sets with 60 cell trajectories. Red and green light inputs are shown in red and green bars, respectively, at the bottom of each panel. Note that each light pattern starts with red light.

### C Parameter inference from experimental data

We demonstrated using simulated data that the diversification of light inputs provides means to increase the information content in the data and thereby allows us to learn model parameters more precisely and with a smaller number of cells.

In this section, we present the results for model calibration using experimental data. We constructed a strain in which the CcaS/CcaR system is driving a fluorescent reporter protein, the repressilator circuit is present but the CcaS/CcaR system is not coupled to it. Cells were growing in a mother machine microfluidic device and a modified overhead projector was used to target light signals at individual cells on the microfluidic chip as shown in Figure 1 in the main text. Experiments to characterise the system were performed with different numbers of cells (12, 30, 60), each of which was assigned to one of six pre-defined different light stimulation inputs (as in the *in silico* study) over a duration of $1-2$ days.

From the raw data, we semi-automatically removed some observed trajectories that obviously could not be used for parameter learning. For example, we removed cells that died during the experiment or those that were not observed correctly due to cell movements. After curation, we were left with 7 - 15 cells per group (G1 - G6), *i.e.* per light signal. From these, in order to parameterise the model, we randomly selected 5 cells from each group or 30 cells in total, visualised in Figure C.14.

Even though the experimental data display similar behaviour to the simulated data (see Figure B.2), the parameter inference reveals additional hurdles that need to be overcome when we deal with the experimental data.

#### C.1 8 parameters are hard to learn jointly

As in our study on simulated data, we use a Metropolis Hastings Markov chain Monte Carlo (MCMC) method based on log-normal proposal distributions with fixed variance for Bayesian inference as described in Appendix B.4. While we can observe a convergence

**Table C.3:** MAP estimators for parameters of the model (11)-(13), obtained using the experimental data set with 30 cell trajectories (Figure C.14) with imposed limits: $n < 10$, $k < 1$, and $c < 15$.

| | $a$ | $b$ | $s$ | $m$ | $h$ | $c$ | $n$ | $k$ |
|---|---|---|---|---|---|---|---|---|
| MAP (8) | 153.2136 | 0.0092 | 3.1098 | 0.0102 | 85.7538 | 10.9291 | 0.02163 | 0.2992 |
| MAP (4) | same | 0.0123 | same | 0.0111 | 1.16e-05 | 5.0901 | same | same |

of log-likelihood values, some parameters do not seem to converge even after a very long run of the MCMC algorithm, while some other parameters show a bimodal behaviour. In particular, the parameters that seem to be most difficult to infer from data are the ones that define the time-dependent rate of the reaction that describes the protein production in the model (B.3), in particular $a, m, c, n$ and $k$.

After imposing bounds on parameters $n < 10$, $k < 1$ and $c < 15$, running a MCMC parameter search for 100 000 iterations, and removing a burn-in period of 40 000 iterations, we still do not observe a clear convergence of the search. Two-dimensional marginals (Figure C.16) show a very strong correlation between some parameters, which makes them very difficult to learn. For example, parameter $a$ shows a strong correlation with parameters $m$, $h$, $c$, $n$ and $k$. Similarly, parameter $m$ shows a strong correlation with $h$, $c$, $n$ and $k$, etc. However, the fitting of the mean using the MAP estimator is still relatively good (Figure C.14).

For this reason, we decide to fix four out of eight parameters: $a$, $n$, $k$ and $s$, to their inferred MAP values (see Table C.3) and run another search for the remaining four parameters: $b$, $m$, $h$ and $c$. Results of this search are given in Figures 7 - 8 and in Table C.3.

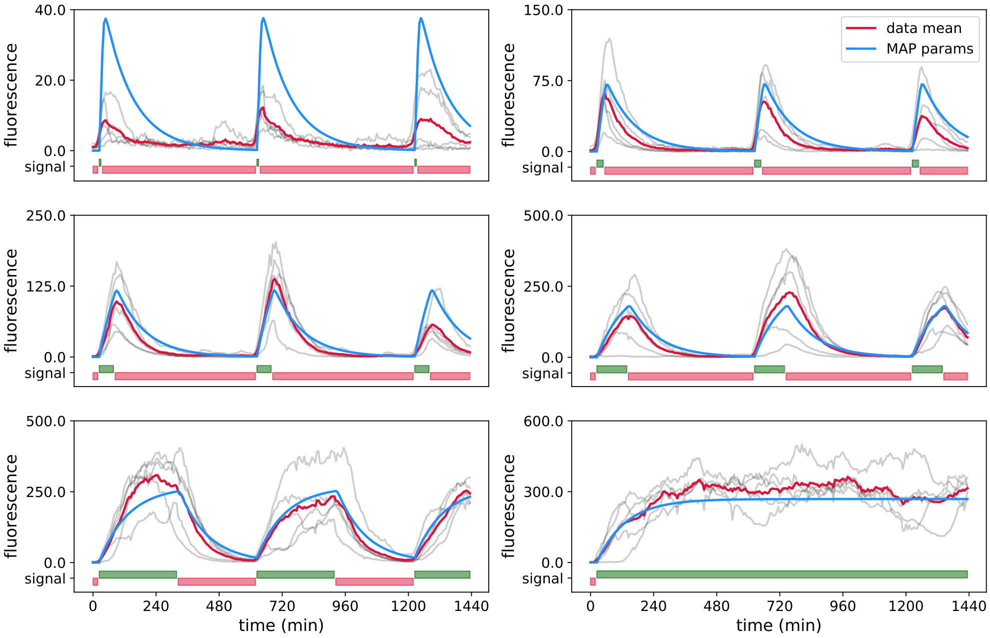


**Figure C.14:** Experimental data, 30 cells in total, 5 cells per light signal. Each trajectory (gray lines) follows one cell over a course of 24 hours, time between every measurement point is 6 min. Red lines are means of data and blue lines ideal means predicted by the model using the MAP estimator of 8 parameters obtained from this same data set, Table C.3. Red and green light inputs are shown in red and green bars, respectively, at the bottom of each panel. Note that each light pattern starts with red light.

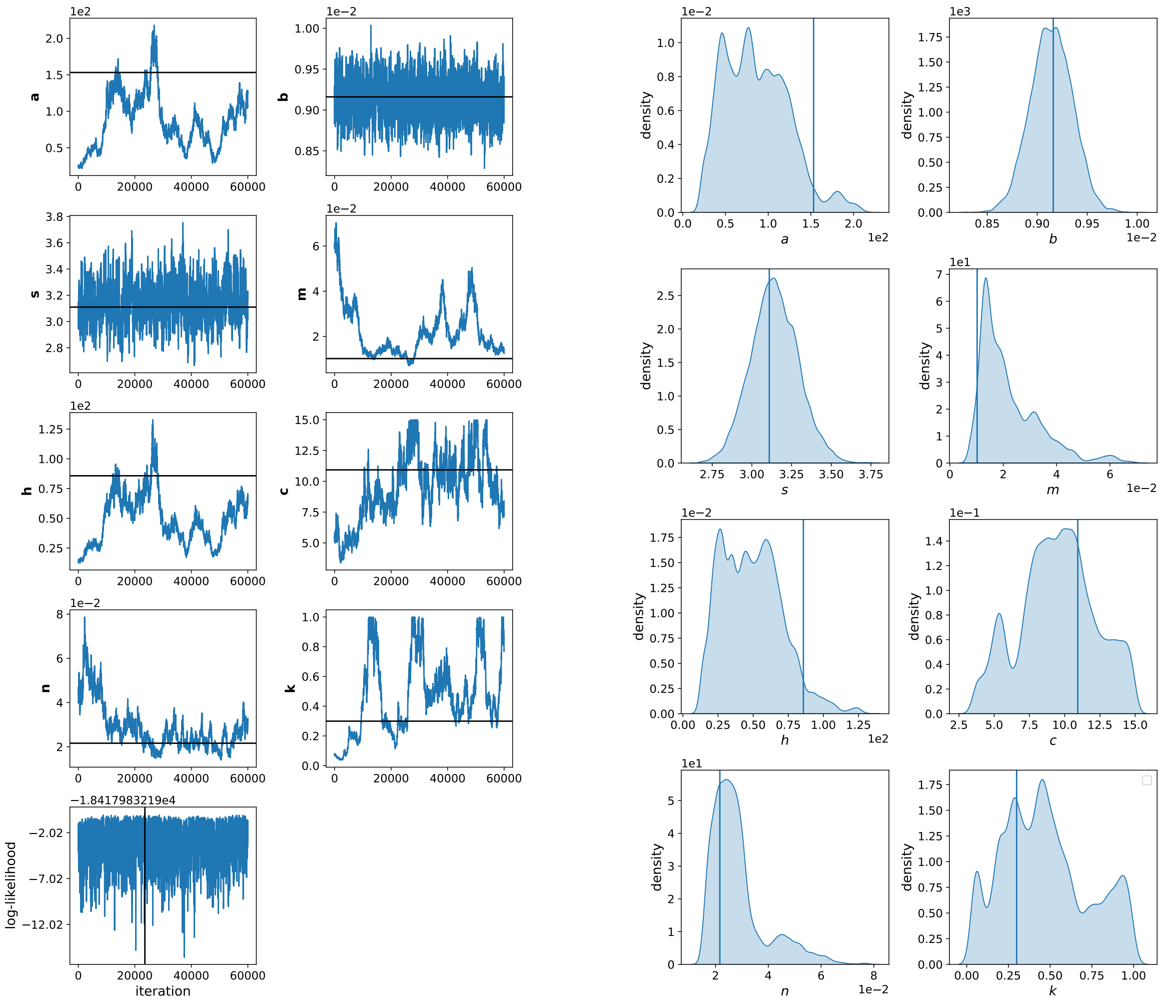


**(a)** MCMC. **(b)** Posteriori distributions.

**Figure C.15: Inference using Algorithm 1 on real data.** **(a)** MCMC chain after a burn-in period of 40 000 iterations, showing the search for each of the parameters and respective values of log-likelihood function. Black lines show the value and the position in the chain of the MAP estimator. **(b)** One-dimensional marginals of posterior distributions. Data used for inference of parameters are 30 cells from the experiments, 5 cells per light group, as in G0, see Figure C.14.

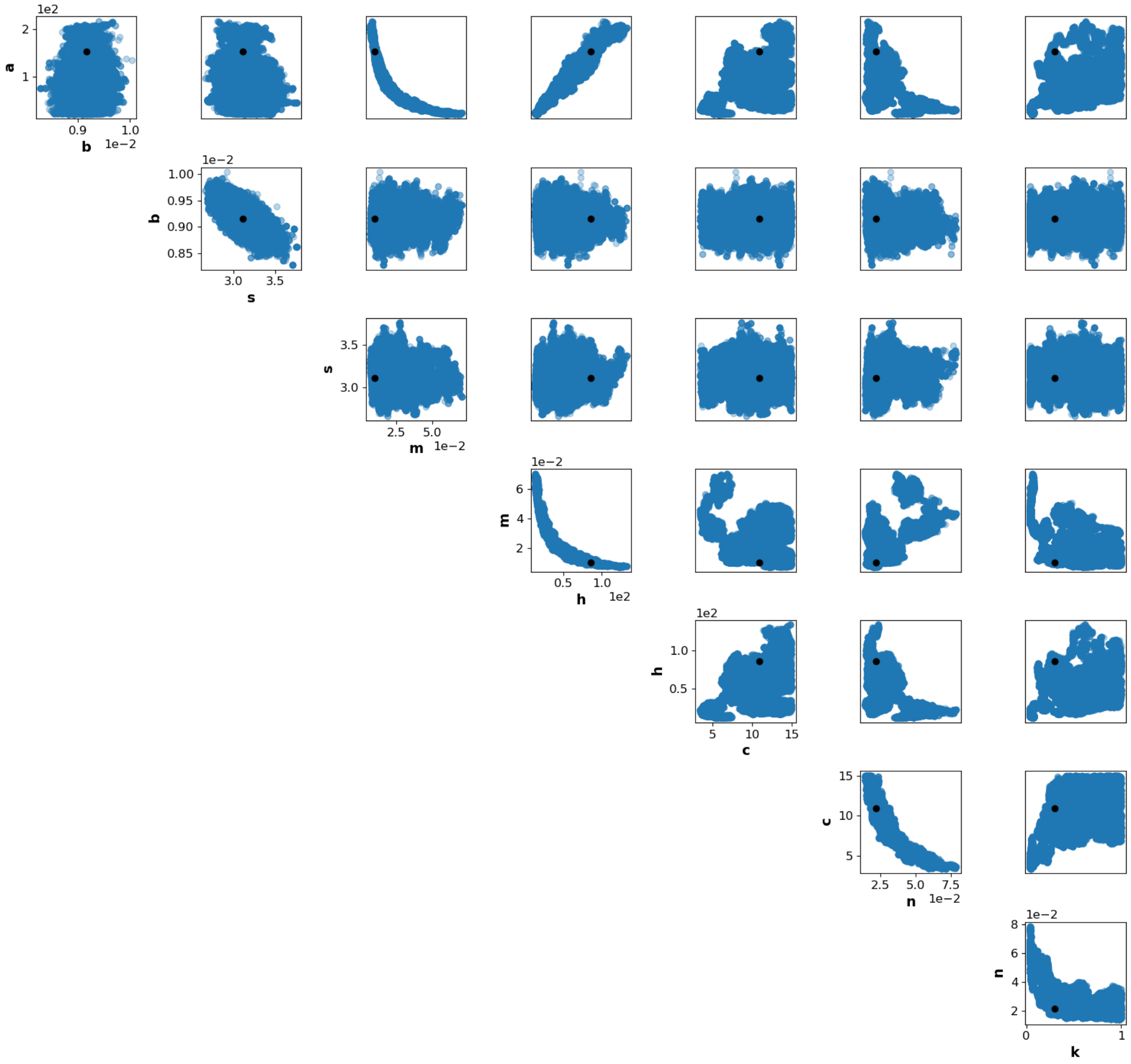


**Figure C.16:** Two-dimensional posterior marginals of the MCMC parameter search using 30 cell trajectories in total, obtained in the experimental setting, 5 cells per light group, as in G0. The MCMC search has 100 000 iterations with a burn-in period of 40 000 iterations. Each iteration takes about 5 seconds of computational time. Black dots show values of the parameters of the MAP estimator.

#### C.2 Bounds on parameters

Here we will try to understand where some of the non-identifiabilities come from and how we chose bounds on the parameters.

##### C.2.1 Bounds on $n$, $k$ and $c$

We take a closer look into the model (B.3) - (B.4). The light activation dynamics is given through the equation

$$\frac{dl(t)}{dt} = u(t) - cl(t). \tag{C.5}$$

We know that $u(t)$ is a piecewise constant function that takes values

$$u(t) = \begin{cases} 0, & t \in [t_{2i}, t_{2i+1}), \quad i = 0, 1, 2, ... \\ 1, & t \in [t_{2i+1}, t_{2i+2}), \quad i = 0, 1, 2, ... \end{cases}$$

where, with the abuse of notation, we denote with $t_i, i = 0, 1, 2, ...$, the times of changes of light signals from green to red and vice versa. Hence, the equation for $l(t)$ has a piecewise solution:

$$l(t) = \begin{cases} l(t_{2i})e^{-c(t-t_{2i})}, & t \in [t_{2i}, t_{2i+1}), \quad i = 0, 1, 2, ... \\ \frac{1}{c} - \left(\frac{1}{c} - l(t_{2i+1})\right) e^{-c(t-t_{2i+1})}, & t \in [t_{2i+1}, t_{2i+2}), \quad i = 0, 1, 2, ... \end{cases} \tag{C.6}$$

With the initial condition $l(t_0) = 0$, it is easy to see that the value for $cl(t)$ is bounded and $0 \leq cl(t) < 1$ for all $t \geq t_0$. Keeping this in mind, we turn to the Hill function,

$$L(t) = \frac{(cl(t))^n}{(cl(t))^n + k^n}. \tag{C.7}$$

One may notice that if $k = 0$, then we have a trivial situation where $L(t) = 1$ for all $t$, in which case there is no effect of light on this system and the value of $n$ does not matter. On the other hand, if $k \gg 1$, since $cl(t) < 1$, $L(t)$ always stays close to 0, and is not significantly affected by the dynamics of the light system. For this reason we impose $0 < k < 1$.

The parameter $n$ is a Hill coefficient that defines the steepness of the function $L(t)$. Theoretically an exponent may take any positive value, practically we will not observe any different shape of the function for big values of $n$. Hence we impose $n < 10$.

Another role of the parameters $c, n$ and $k$ is to determine the maximum value of $L(t)$ and therefore the maximum value of the rate $aE(t)L(t)$. If we take a look at our data we notice that short green light signals are always followed with long red light signals. Thanks to that we may assume that $L(t) = 0$ at the beginning of the subinterval where the green signal, $u(t) = 1$, is applied; and the maximum value, $L_{max}$, is reached at the end of that subinterval. We derive $L_{max}$ from (C.7) and (C.6), using $l(t_{2i+1}) = 0$, to be

$$L_{max} = \frac{1}{1 + \left(\frac{k}{1-e^{-cT}}\right)^n}, \tag{C.8}$$

where $T$ is the length of a green light signal, which varies in the experiment from 12 to 1440 min. Examining the behaviour of this function, taking values of $T$ from 12 to 1440, values of $k$ from 0 to 1, and $n$ from 0 to 10 and observing the dependence of $L_{max}$ with respect to $c$, we conclude that in all cases $L_{max}(c)$ is virtually constant for $c > 2$. When we take $T = 1$, $L_{max}(c)$ is virtually constant for $c > 10$.

One may also notice that every time $E(t) = 0$, the time-dependent rate $aE(t)L(t)$ of production of the fluorescent protein $F$ becomes zero. During that time the parameters of $L(t)$ are not relevant. If we have a presence of $E(t)$ almost continuously (which would reflect in a high value of parameter $m$) this might not be an issue. But a small value of the parameter $m$ means that there are many periods when $E(t) = 0$ and hence we have reduced time to learn parameters of $L(t)$. However, since our algorithm works with approximation of the average behaviour and not the exact model, the value that really determines the dynamics of the system is the product $a \cdot m$, and we would still be able to say something about the system if this product is not very small.

In order to control somewhat possible pitfalls of the parameter inference we use the bounds on parameters $n < 10$, $k < 1$ and $c < 15$. While these bounds are sufficient to learn parameters from simulated data, we still have issues of convergence when using experimental data, as described in the previous section. This may be related to the model of the measurement noise: we assume an additive Gaussian measurement noise (see Section 2, eq. (3)), but we do not know if this is a good assumption for the experimental data. Further investigation in data noise levels would be needed.

##### C.2.2 Switch times and parameter c

The Hill function, $L(t)$, is a switch function, although continuous and differentiable, that switches between 0 and some maximal value $L_{max}$. We define *a switch time*, $t'$, to be a moment in which $cl(t') = k$ and $L(t') = \frac{1}{2}$. In the limit case when $n \to \infty$, $t'$ would be a point of discontinuity, because $cl(t) > k$ would give $L(t) = 0$ and $cl(t) < k$ would give $L(t) = 1$. In order to understand how the switch time depends on the parameter $c$, we use the expression for $l(t)$ given in (C.6), to find

$$t' = \begin{cases} t_{2i} + \frac{1}{c}\ln\left(\frac{cl(t_{2i})}{k}\right), & \text{for } u(t) = 0, \quad t \in [t_{2i}, t_{2i+1}), \quad i = 0, 1, 2, ... \\ t_{2i+1} + \frac{1}{c}\ln\left(\frac{cl(t_{2i+1})-1}{k-1}\right), & \text{for } u(t) = 1, \quad t \in [t_{2i+1}, t_{2i+2}), \quad i = 0, 1, 2, ... \end{cases} \tag{C.9}$$

Notice that a switch may happen on every subinterval where light signal, $u(t)$, is constant and it depends on the initial value of $l(t)$ on the given subinterval. For simplicity, in what follows we denote the initial value of $l(t)$ on the given subinterval with $l_0$. On the subintervals where $u(t) = 0$, the function $L(t)$ is decreasing and it may switch after some time

$$\delta_{\text{fall}} = \frac{1}{c}\ln\left(\frac{cl_0}{k}\right). \tag{C.10}$$

The profile of this function looks as follows: for $c$ very small such that $cl_0 < k$ we have $\delta_{\text{fall}}(c) < 0$, which is not a relevant part of the curve; the curve crosses 0 when $cl_0 = k$; reaches its maximum value $\delta_{\text{fall}}(c) = 1/c$ when $cl_0 = ek$; and then falls back asymptotically to 0 for large $c$, with a rather flat asymptotic tail. As one may notice, the function $\delta_{\text{fall}}(c)$ is not monotone. This implies that two or more values of parameter $c$ may explain equally well the switch time of the fall. On the subintervals where $u(t) = 1$, the function $L(t)$ is increasing and it may switch after some time

$$\delta_{\text{rise}} = \frac{1}{c}\ln\left(\frac{cl_0 - 1}{k - 1}\right). \tag{C.11}$$

The profile of this function looks as follows: if $c \to 0$ then $\delta_{\text{rise}}(c) \to +\infty$; it is monotonically decreasing and hits 0 when $cl_0 = k$; for $c$ big such that $cl_0 > k$ we have $\delta_{\text{rise}} < 0$, which is not a relevant part of the curve. In this case we assume that $l_0$ is rather small and hence $c$ might be rather large. Theoretically, we should be able to learn the parameter $c$ from

$\delta_{\text{rise}}$, if $\delta_{\text{rise}}$ is observed in data, *i.e.* if the green light signal is long enough. However, if the algorithm suggests too large values for $c$ the switch times $\delta_{\text{rise}}$ and $\delta_{\text{fall}}$ would be so small that we would not be able to observe them in our data with time resolution of 6 min. Notice, however, that we expressed only first order approximations for $\delta_{\text{fall}}$ and $\delta_{\text{rise}}$ as the initial values $l_0$ also depend on $c$.
